# Control of gene output by intron RNA structure

**DOI:** 10.1101/2025.11.08.687378

**Authors:** Leonard Schärfen, Pernille Bech, Paulina Podszywałow-Bartnicka, Karla M. Neugebauer

## Abstract

During mRNA biogenesis, RNA folding can promote or antagonize transcript processing. The effects of intron structure on mRNA and protein levels remain largely unexplored, although introns account for the bulk of nascent RNA. Here we systematically probe the effect of intron structure using massively parallel reporter assays. We show that base pairing modulates gene expression across orders of magnitude through inhibitory RNA structures at splice sites and at newly identified regions. Conversely, poor splicing of human β-globin pre-mRNA could be improved by sequence alterations that reduce base pairing. For large libraries of RNA structures differing in stability, machine learning models could nearly fully explain observed gene output. Structure destabilizing mutations emerge rapidly under selective pressure. Thus, formation of RNA structures as dictated by intron sequence provides a simple, powerful means to adjust gene expression.

---

RNA structure formation, driven by intramolecular base pairing, is a feature of every transcript. Because folding occurs during transcription and within the time frame of pre-mRNA splicing (*1–4*), we sought to ask how RNA structure contributes to splicing regulation. Indeed, our understanding of the mechanistic control of splicing efficiency and rate remains frustratingly obscure. The interplay between the number and length of introns and exons, splicing enhancer and silencer sequences, and hundreds of RNA binding proteins renders splicing efficiency highly context-dependent and locus-specific (*5–7*). This complexity has raised the question of why eukaryotic genes even have introns that occupy up to 90% of protein coding gene space in mammals (*8*). While high-resolution structural models of the spliceosome at different stages of the splicing reaction are available (*9*), the structure of the RNA substrate is usually not resolved in CryoEM maps and rarely considered. Despite the ubiquity of RNA structure formation, only select examples of structures regulating splicing have been reported (*10–14*). Progress in detecting intron base pairing patterns has been made (*1, 2, 4, 15*), but there are many limitations to identifying structures and determining their functions in short-lived pre-mRNAs. Because a quantitative description of splicing regulation through intron folding is missing, we lack approaches that could search for or predict the stabilities and locations of intronic structures with the capacity to regulate splicing.

Splicing reporters can address gaps in our understanding of dynamic RNA folding and its impacts on gene expression, because targeted changes to isolated regulatory elements can be separated from the remaining sequence. Previously, studies investigating alternative splicing have been carried out by interrogating exon sequence variants using RNAseq (*16– 21*). However, the study of intron sequence poses a unique challenge for high-throughput approaches, because the intron is removed and degraded, requiring barcodes and deep sequencing for detection (*22, 23*). Here we systematically address the role of structure across entire introns, including evolutionarily young and ancient features, using *Saccharomyces cerevisiae*. Budding yeast is an excellent model system, because previous studies have already shown that endogenous introns placed into reporter genes exhibit a wide range of expression levels, showing that intron sequence affects splicing efficiency (*24*). Several examples have shown that splicing can be modulated by RNA structure, suggesting that accessibility to essential regulatory elements like splice sites is crucial (*13, 14, 25*). However, sequence analysis alone is currently insufficient to identify such structures and their effect on splicing, necessitating the massively parallel experimental approach we pursue here. We derive rules on the location and stability of RNA structures, their impacts on splicing and relationship to gene output and, remarkably, can replicate these regulatory effects in mammalian cells. Finally, we show that intronic mutations that alter RNA structure are likely to emerge during natural selection.

## Results

### High precision measurement of gene output

We sought to develop a reporter assay that allows quantification of protein production with high precision and resolution in genetic backgrounds differing only in intron sequence. Both nascent RNA folding and pre-mRNA splicing are fundamental to all eukaryotes, including the yeast *S. cerevisiae*, with highly conserved core spliceosomal components (*26*). We therefore constructed a reporter gene for recombinase-mediated integration into the *S. cerevisiae* genome, which codes for yeast-optimized (*27*) and HA-tagged mNeongreen (mNeon). This fluorescent protein can only be produced when the single intron is spliced, because it is otherwise out of frame (*28, 29*). The short 85 nt first exon matches the naturally occurring length of first exons in yeast and transcription is driven by the strong endogenous *TDH3* promoter (Fig. 1a). We measured mNeon levels at the single cell level using flow cytometry concurrent with red fluorescence from the mScarletI (*27*) (mScar) protein, driven by the same expression platform at a different genomic locus (Fig. 1b). By calculating the log ratio of green and red fluorescence for each cell, splicing of the reporter is separated from general extrinsic gene expression noise (*30*) caused by cell size, age, cycle, etc. This allows precise quantification of relative protein production by normalizing the log ratio such that it lies between the negative (no mNeon) and positive (no splicing required; intronless) control (Fig. 1c). We refer to this normalized value as “gene output”. This is a protein-level proxy that integrates splicing efficiency with mRNA stability and translation; however, the mRNA sequence is the same after processing. Thus, splicing is expected to be the major contributor to this measurement. As reported previously (*24*), different endogenous yeast introns negatively affected gene output, while others had no effect or slightly increased gene output relative to the intronless control (Fig. 1c).

**Fig. 1:**
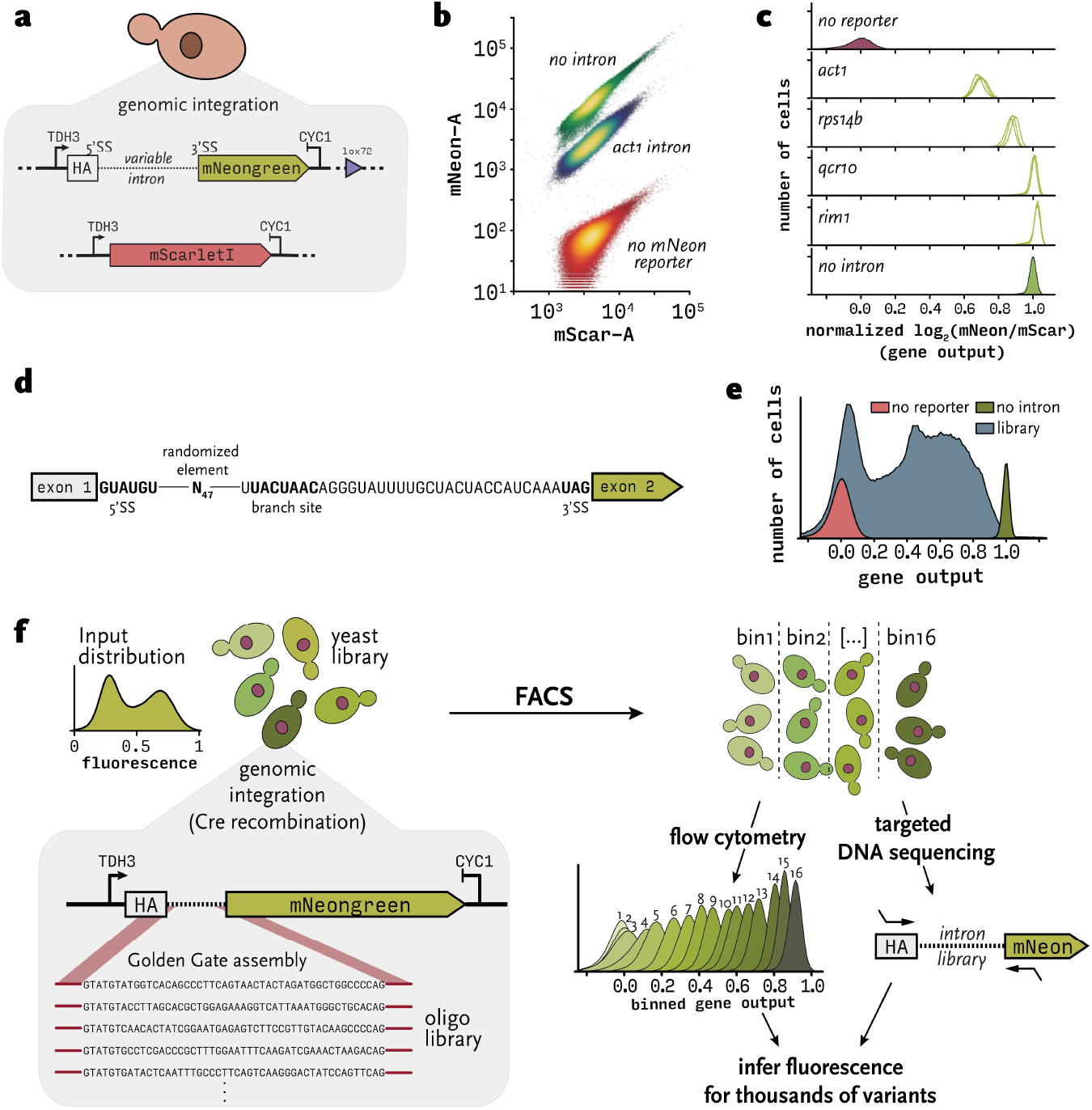
Precise measurement of reporter gene output for thousands of intron variants. **(a)** Two genes coding for fluorescent proteins are integrated at different loci into the genome of *S. cerevisiae*. The splicing reporter with HA as exon 1, a modular intron, and mNeongreen is integrated at the *MET17* locus via a Cre/*lox* landing pad. mScarletI is driven by the same expression platform at the *HIS3* locus. **(b)** Flow cytometry measurement of fluorescent protein amount expressed from reporter genes. 10^6^ cells shown for each strain. **(c)** Gene output values for different endogenous introns. For each cell, splicing efficiency is calculated as the log2 of green over red fluorescence and then normalized such that a value of zero corresponds to no expression of mNeongreen, and a value of one corresponds to expression without the need for splicing. Three biological replicates are plotted for each intron. **(d)** Design of a partially randomized *BOS1* intron. **(e)** Fluorescence distribution of partially randomized intron library. **(f)** Sort-seq starts with high-throughput genomic integration of an intron variant library. Cells are then sorted according to gene output into 16 bins of similar size. Cultures from each bin are again subjected to flow cytometry to determine the output fluorescence distribution of each bin (output distribution for all 16 bins of the randomized library in (d) is shown). The variable region is amplified and sequenced on the Illumina platform for each bin.

### Randomized libraries reveal intron RNA structures that modulate gene output

Effect size characterization of RNA structure-based splicing regulation requires connecting structure formation with gene output for many thousands of sequence variants. We adapted the reporter assay described above to a high-throughput format. First, we generated a reporter gene library with a random N_47_ element based on the endogenous *BOS1* intron, randomizing sequence between 5’SS and branch site (Fig. 1d). This design ensures that variation in splicing efficiency is not due to altered splice sites and/or the short sequence between branch site and 3’SS, which were previously shown to affect splicing (*31–33*). The intron library was integrated into the genome of *S. cerevisiae* via Cre recombination with a landing pad locus and selected with hygromycin (see Methods), ensuring that every cell harbors one copy of the reporter gene. In total, we sampled approximately 10^5^ colonies (each corresponding to one unique variant) from the 10^28^ possible sequences that this library design permits. When subjected to flow cytometry, splicing efficiencies spanned almost the full range of detection, showing that intron sequence strongly regulates splicing independent of 5’SS or the 3’-end of the intron (Fig. 1e). To assign an accurate gene output value to each intron sequence, we performed Sort-seq (*34, 35*) (Fig. 1f): We used FACS to sort the yeast library into 16 overlapping bins according to gene output, grew the sorted cultures, and again performed flow cytometry to measure output distributions. We next isolated genomic DNA from each bin, amplified the intron region and sequenced on the Illumina platform. We extracted read counts for each variant and calculated the fraction of reads found in each bin (Fig. S1a). As expected, read counts from neighboring but not from distant bins were highly correlated (Fig. S1b). Normalized read counts for each variant combined with the exact position of each bin allowed accurate estimation of the original fluorescence value (Fig. S1c, see Methods). Only variants detected in at least two bins were retained, resulting in accurate gene output measurements for 90,064 variants.

This rich dataset enabled us to associate the structural landscape of the library with gene output. To test whether gene output depends on RNA structure formation in this system, we calculated the ensemble free energy (EFE) using ViennaRNA (*36*) for each random intron; the EFE signifies the free energy of all possible secondary structures that each RNA can adopt, weighted by their probabilities. We first asked if we could determine the optimal sequence window for structure prediction by comparing correlation coefficients for EFE and the measured gene output for every combination of exon and intron coordinates (Fig. 2a). This approach surveys different lengths of RNA, exploring sequence space spanning from the first exon through the entire intron and into to the second exon. The resulting correlation coefficient matrix shows the correlation between EFE and gene output sharply increases when >35nt of 1^st^ exon (upstream of 5’SS) and >60 of intron (downstream of 5’SS) are considered. The strong positive correlation between EFE and gene output at the optimal prediction window confirms that RNA structure formation is a major regulator of splicing (Fig. 2b).

**Fig. 2:**
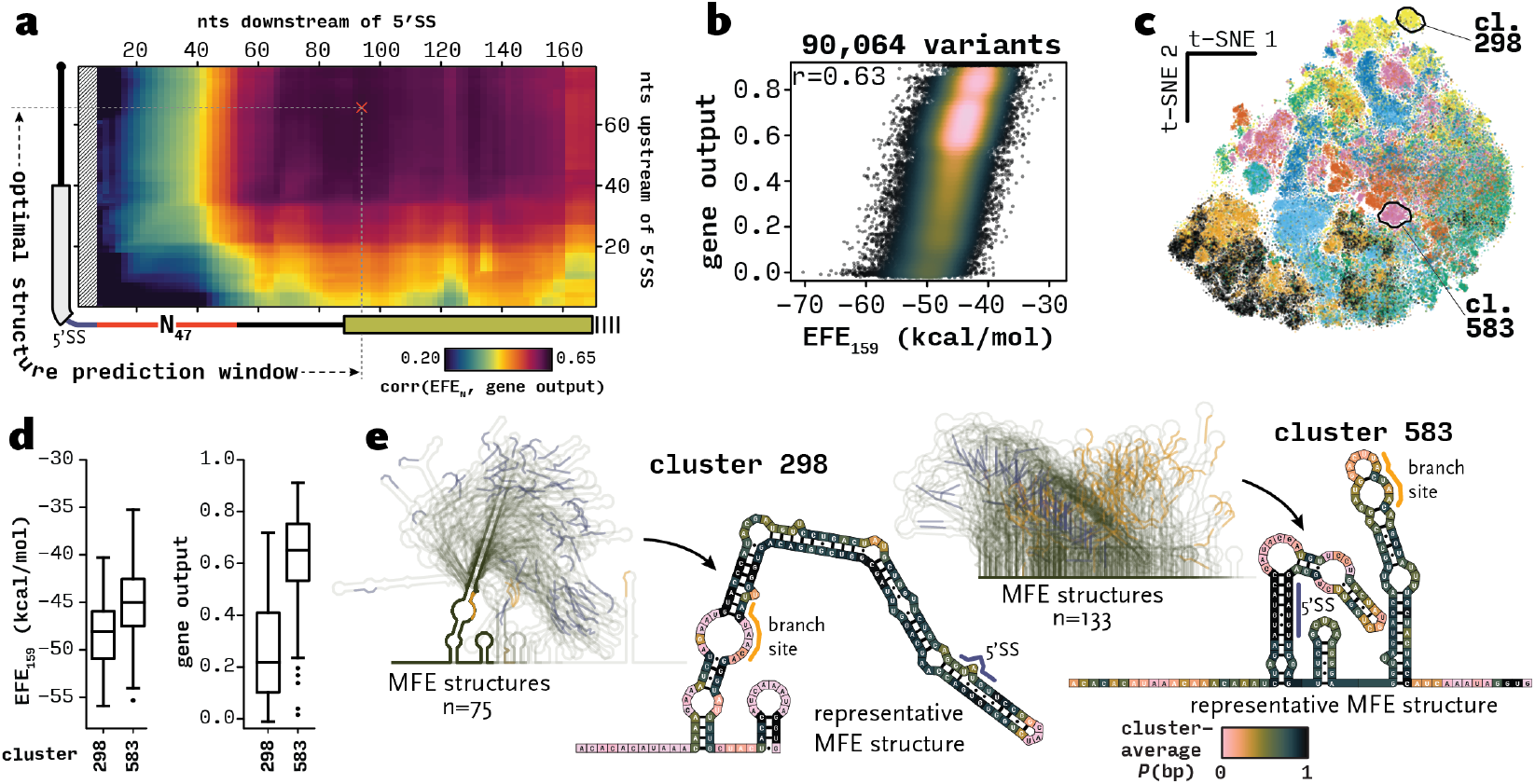
Predicted RNA structures for randomized introns explain gene output only partially. **(a)** Pearson correlation coefficients of EFE and gene output for varying structure prediction windows. Y-axis represents included sequence upstream of the 5’SS for prediction and the x-axis represents included sequence downstream. Red N_47_ signifies the variable sequence between 5’SS and branch site (see Fig. 1d). **(b)** EFE versus gene output for the optimal window size, colored according to density. r: Pearson correlation coefficient. **(c)** t-SNE embedding of the positional entropy matrix, colored by clusters obtained from hierarchical clustering of the *P*(bp) matrix. Clusters analyzed in (d) and (e) are highlighted. **(d)** EFE and gene output distributions for clusters 298 and 583. **(e)** Overlayed minimum free energy (MFE) structures of all member sequences in clusters 298 and 583. 5’SS and branch site are highlighted in blue and orange, respectively. The MFE structures best described by average *P*(bp) across each cluster are shown in detail, colored by cluster average *P*(bp).

To approach a mechanistic model for how RNA structure regulates splicing, an understanding of effects at the single nucleotide level is required. For example, a sequence might fold into structures with an overall high free energy that may only correlate with function depending on the extent of base pairing with a functional site, like a 5’SS, branch site, or unknown functional site. We therefore calculated *pairing probabilities P*(bp), i.e., the weighted fraction of paired nucleotides across all possible structures in the ensemble at each position, as well as *positional entropy*, i.e., the uncertainty or variability in structural state of each position, for every nucleotide of each random sequence within the optimal window. We embedded the 159-by-90,064 entropy matrix into two dimensions using t-SNE and observed distinct clusters that were well separated by hierarchical clustering of the *P*(bp) matrix (Fig. 2c, Fig. S1d). Each cluster represents a set of sequences that are structurally similar. If structure is a major determinant of splicing, we would expect each cluster to exhibit a distinct gene output distribution measured by Sort-seq. By arranging clusters by standard deviation of their gene output values, we found select clusters with relatively narrow distributions (Fig. S1e). For example, cluster 298 is characterized by low EFE and low gene output (Fig. 2d), predicted to form a single stable stem loop that sequesters both 5’SS and branch site (Fig. 2e). In contrast, cluster 583 is predicted to be less stable with sequestered 5’SS but free branch site. Despite these examples, most clusters exhibit broad gene output distributions. We were also unable to train random forest regressors or convolutional neural networks to predict gene output from pairing probabilities or from sequence to perform better than a linear EFE predictor (Fig. 2b), in agreement with previous studies (*33, 37*). Most likely, the comparatively small sample (10^5^) from a large sequence space (10^28^) does not contain enough structurally similar variants to inform effective models. This implies that smaller structural variations than those capturable in a random library are needed to address biological relevance.

### A “walking stem” strategy scans intron space for RNA structure-based regulation

To devise general rules of intronic structure formation and its impact on splicing, we carefully designed variant libraries that address RNA structure directly. Specifically, what elements of an intron are subject to splicing inhibition by intramolecular base pairing? We developed a generalizable library design strategy that tests the effect of intramolecular base pairing at every possible position of functional sequence (Fig. 3a). We chose the 82 nt intron of the *GOT1* gene as a basis for library design, because it is predicted to be unstructured (*10*), matches the average length for introns in non-ribosomal protein genes, and the wildtype sequence did not decrease protein production in the reporter gene context (Fig. 1c). A 19 nt artificial sequence was inserted at *each possible* position, similar to two previous reports where sequences were inserted at one individual position (*38, 39*). Using the RNA design algorithm antaRNA (*40*), the inserted sequence was designed to form a 13-base pair stem capped by a tetraloop, sequestering either the upstream or downstream sequence (relative to the position of insertion) in base pairing interactions. To control for sequence-specific effects that might obscure the signal caused by sequestering insertions, three distinct stem loops were generated per position, and supplemented with destabilized versions of each by installing three point mutations that maximally increase free energy of the target structure (*41*). We also included a set of sequences specifically designed to remain single-stranded (Fig. S2a). Altogether, the library features 6 inserts forming stem loops, 12 destabilized stem loops, and 10 unstructured insertions per position for a total of 1,998 intron variants. The predicted *P*(bp) matrices reproduced the expected patterns of RNA folding, with fully formed stem loop (Fig. 3b), destabilized stem loop or single-stranded sequence (Fig. S2b) “walking” across the intron.

**Fig. 3:**
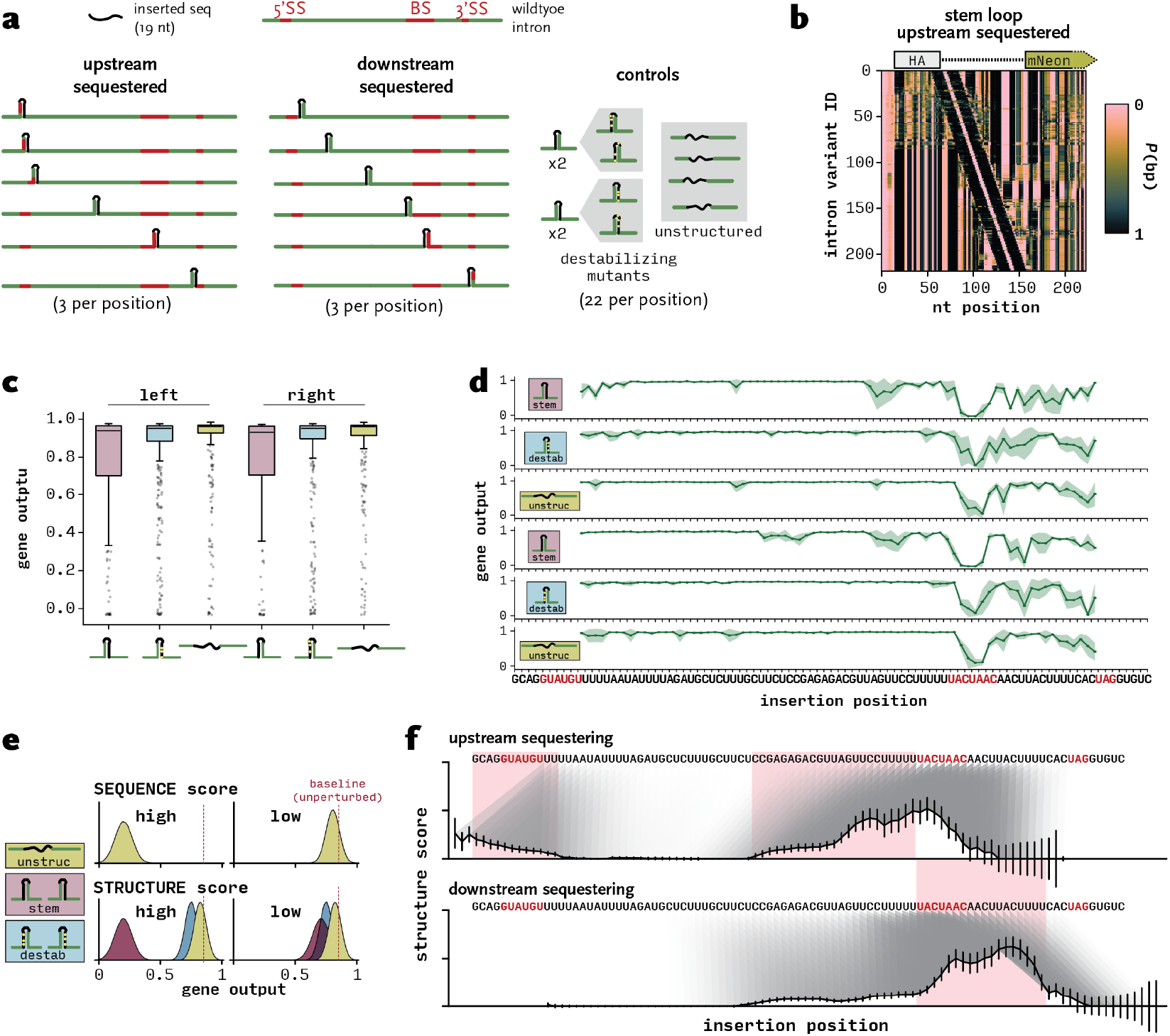
Systematic discovery of intron elements that are sensitive to base pairing. **(a)** Library design principle. A modular 19 nt sequence is inserted into the target sequence at every position. The insert is algorithmically designed to pair with upstream or downstream sequence. Each designed insert is accompanied by control inserts that harbor three point mutations to destabilize the target structure, as well as inserts specifically designed to *not* pair with surrounding sequence. **(b)** Predicted pairing probabilities for each nucleotide of the designed *GOT1* intron variants harboring the insert sequestering upstream sequence. Each row of the matrix represents one unique intron. **(c)** Observed gene output distributions determined by Sort-seq for each type of insert. **(d)** Average splicing efficiency per insertion position for each insert type. Mean ± sd from all variants of the same insert type per position. **(e)** Schematic showing how structure score and sequence score in (f) are interpreted. **(f)** Structure score calculated from all insert types at each position. Identified structure-sensitive elements are highlighted in pink.

We synthesized and cloned the library, integrated it into *S. cerevisiae* and performed Sort-seq as described above. Overall, inserts designed to form stable stem loops had the largest effect on gene output (Fig. 3c). Breaking down the types of inserts along the intron, each construct’s positional effect can be visualized (Fig. 3d). As expected, any insertion disrupted gene output when placed inside the branch site. The region between branch site and 3’SS was also affected by every insertion, likely because it is subject to distance and sequence constraints (*42*). Taken together, these observations suggest that insert-mediated structure formation decreases gene output by impeding splicing.

Upstream-sequestering insertions disrupted splicing when close to the 5’-end of the intron, suggesting that sequestration of the 5’SS inhibits association of U1 snRNP (Fig. 3d). Downstream-sequestering, destabilized and single-stranded insertions at the same position had little to no effect, confirming that the decrease in splicing efficiency is structure and not sequence dependent. Intriguingly, a similar effect was observed upstream of the branch site for downstream-sequestering variants, indicating that U2 snRNP association was also inhibited by sequestered branch site sequences. Upstream-sequestering insertions just upstream of the branch site decreased splicing efficiency slightly, and this result is complemented by downstream-sequestering variants at the opposite side of this sequence, indicating that base pairing of sequence upstream of the branch site is unfavorable (Fig. 3d). To systematically distinguish effects of inserted structures from the effects of disrupting a required sequence, such as a splice site, we first derived a “sequence score” that describes insertion effects that also occurred when destabilized structures and unstructured sequence was inserted. We then extended this scoring system to summarize the observed effects from all sub-libraries (Fig. 3e) where this “structure score” is high when the decrease in gene output caused by the insertion is large relative to the sequence score. Importantly, the score flagged the 5’SS, branch site, and also a novel element upstream of the branch site (Fig. 3f). This analysis also highlights the striking flexibility of introns to accommodate sequence insertions, including structured inserts, at most positions without impeding splicing efficiency at all. Since we systematically accounted for sequence-based effects, this experiment establishes that RNA structure constrains efficient splicing at the substrate level.

### RNA structure stability fine-tunes gene output

The discovery of structure-sensitive intronic elements raises several new questions. First, the biochemistry of how snRNPs compete with base-paired splice sites is unclear. There might exist a mechanism to actively disrupt base pairing, e.g. through helicase activity. In this case, the stability of the intramolecular stem might not matter. Conversely, stability would be important if RNA components of snRNPs access splice sites through strand invasion and displacement of intramolecular pairs. Modulating structure stability could thereby substantially impact splicing efficiency and gene output.

To systematically test how structure stability across structure-sensitive sites affects splicing, we designed a set of combinatorial libraries (Fig. 4a) to be assayed with the Sort-seq splicing reporter. Analogous to the previous experiment, we designed a short insert sequence to sequester each splice site in base pairing interactions. To introduce sequence diversity, libraries were created starting from a single oligonucleotide that was synthesized with degenerate bases. While the nucleotides making up the bottom 8-10 base-pairs of the stem are invariable, degenerate bases placed opposite of the splice site essentially randomize (or titrate) availability of the functional sequence (Fig. 4b). Each combinatorial oligonucleotide library was cloned such that all variants were represented in the resulting yeast library (see Methods; Table S1). Sort-seq was then performed as described above, yielding gene output values for each variant, which showed good agreement with individually measured clones (Fig. S3a). We again found the optimal window for structure prediction using a scanning approach for each library (Fig. S3b). Like the fully randomized library, gene output of 5’SS and branch site variants exhibited strong correlations with intron structure stability, albeit in a clearly non-linear manner (Fig. 4c). Interestingly, splicing efficiency values for introns containing a stem loop covering the 3’SS showed a weak negative correlation with EFE, perhaps due to the different mechanism of recognition compared to 5’SS and branch site (see Discussion). To test reproducibility and to ensure that we measured structure and not sequence-specific effects, we generated another iteration of the 5’SS library (Fig. 2b alternative nucleotides, Tab. S1) and cloned it into reporter genes with HA-tag as exon 1 (which the intron is designed to pair with), as well as MYC-tag as exon 1 (no specific pairing) and performed a separate Sort-seq experiment (Fig. 4d). While this library in the HA context follows the same trend as the previous one (Fig. 2c), the same library in the MYC context displays almost exclusively high fluorescence, strongly suggesting that base pairing (rather than other effects) is the reason for reduction in gene output.

**Fig. 4:**
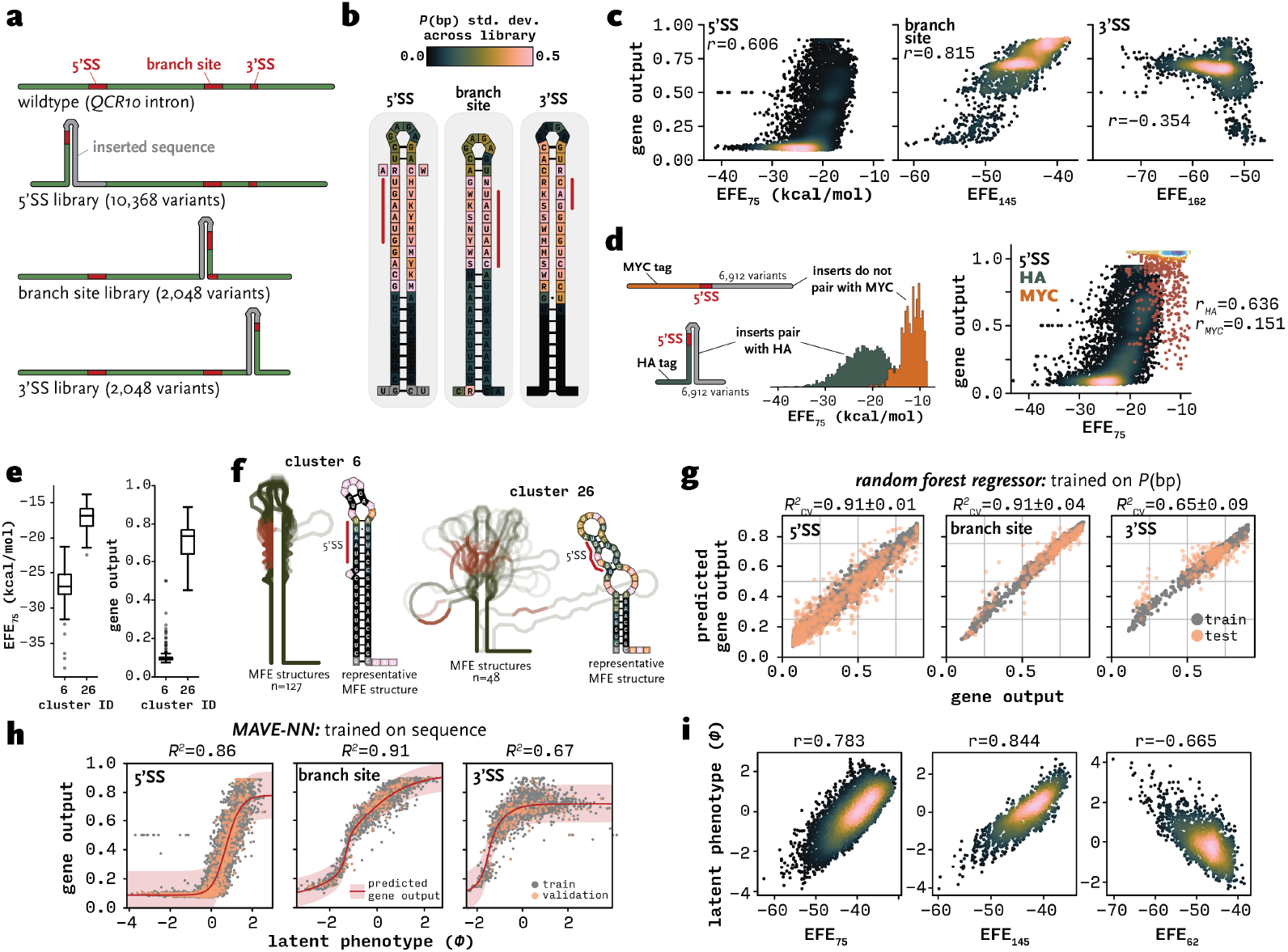
Combinatorial structure libraries reveal highly sensitive response of gene output to stem loop stability. **(a)** Library design principle. An artificially designed sequence is inserted into the target intron (*QCR10* gene) in the vicinity of splice sites, such that the site is sequestered in base pairing interactions. The sequence opposite of the splice site is partially randomized, yielding libraries that represent a range of base pairing states. **(b)** Detailed models of fully formed stem loops differentially sequestering the three splice sites, colored by variability in pairing probability (standard deviation) across each library. **(c)** Relationship of predicted EFE and gene output for each intron variant. *r*: Pearson correlation coefficient. **(d)** EFE distributions and relationship to gene output for a second iteration of the 5’SS stem loop library in the HA and MYC context for exon 1. **(e)** EFE and gene output distributions for clusters 6 and 26. **(f)** Overlayed minimum free energy (MFE) structures of all member sequences in clusters 6 and 26. 5’SS is highlighted in red. The MFE structures best described by average *P*(bp) across each cluster are shown in detail, colored by cluster average *P*(bp) according to color bar in (d). **(g)** Predictive performance of random forest model trained to predict gene output only from pairing probabilities. *R*^2^: coefficient of determination calculated using 30-fold cross-validation. **(h)** Neural network-based inference of interpretable latent phenotype from sequence only. Global epistasis regression model with Gaussian noise and pairwise interactions trained using MAVE-NN with 10% test and 10% validation data. *R*^2^: coefficient of determination calculated using validation data. **(i)** Relationship of predicted EFE to latent phenotype inferred from MAVE-NN model trained using sequence only. *r*: Pearson correlation coefficient.

To compare the information content inherent to this combinatorial approach with the fully random library, we proceeded to calculate *P*(bp) for each variant (Fig. S3c) and clustered them using hierarchical clustering (Fig. S3d). In contrast to the random library, many clusters with low variability in gene output emerged (Fig. S3e). For example, cluster 6 exhibited extremely low fluorescence and low EFE, consistent with a stable stem loop with sequestered 5’SS (Fig. 4e, f). Conversely, cluster 26 contained introns with less stable stem loops, exposed 5’SS and therefore high average fluorescence. This indicates that *P*(bp) is a strong predictor of function for these libraries.

### Machine learning predicts gene output from intron structure or sequence

Predicting phenotype from genotype is one of the central goals in genome biology. To predict gene output from nucleotide sequence in the context of the intron variants described above, we can use knowledge-based models of how RNA folds to extract a latent phenotype (*P*(bp) or EFE) from the genotype. We trained random forest regression models to predict splicing efficiency from the latent phenotype, i.e. from *P*(bp) (Fig. 4g, Fig. S3f). Models for 5’SS and branch site performed exceptionally well, indicating that 1) our thermodynamic models of RNA folding represent reality accurately – even in the cellular context – and 2) RNA structure is the major regulator of splicing in this reporter system.

The 3’SS model accounts for most of the variance in the data, but due to reasons discussed below, RNA structure is likely not the only contributor to splicing outcome for these variants. Random forest regression models allow us to query how much each feature contributes to the prediction. Nucleotides directly adjacent to the splice sites, as well as the designed stem in general have the most predictive power (Fig. S3g). Surprisingly, neither the splice sites themselves nor specifically the variable nucleotides were among the most important positions, perhaps indicating that overall stem loop stability rather than exposed splice site nucleotides are important for enabling recognition. As a negative control, models trained on the 5’SS library in the MYC context had little predictive power (Fig. S4a). Models trained on separate 5’SS libraries in the HA context had predictive power on their respective counterpart that had been independently measured, despite representing different sets of stem loop variants, suggesting that models are not overfitting (Fig. S4b).

As an alternative to explicitly incorporating thermodynamic RNA structure predictions into models, we can attempt to derive a latent phenotype *de novo* by training a model purely on sequence, without prior knowledge. Such models have previously been developed specifically for handling massively parallel reporter assay data. We used MAVE-NN,(*43*) which infers genotype-phenotype maps through a specialized neural network architecture, to train regression models with only variable nucleotides as one-hot encoded features. These models performed similarly to the random forest regressors with strong predictive power (Fig. 4h). The neural network architecture allows us to access the latent phenotype, which represents an optimal embedding of multiple predictors. Strikingly, the inferred latent phenotype correlated well with EFE (Fig. 4i), indicating that the model learns a representation of RNA structure to make gene output predictions.

### RNA structure-based tuning of β-globin splicing in mammalian cells

Splicing regulation is more complex in mammalian cells, where degenerate splice sites need to be identified by a host of accessory proteins, such as helicases and RNA binding proteins that can stabilize single stranded RNA (*5, 6*). These features could counteract the direct relationship between stem loop stability and gene output than we have seen in the yeast system. To test the generality of our findings, we constructed an analogous dual-color reporter plasmid (pDual) for transfection into tissue culture cells (Fig. 5a). As in the yeast system, mScarletI (mScar) was expressed to control for extrinsic noise. The mammalian reporter gene features the 1^st^ exon and the intron of the human β-globin (*HBB*) gene, and mNeongreen (mNeon) as exon 2. Interestingly, human *HBB* pre-mRNA is known to be inefficiently spliced (*44*) and is predicted to form a stem loop across the exon-intron junction (Fig. S5a). As in the yeast system, splicing was required to produce mNeon protein. We transiently transfected the pDual plasmid containing the wildtype intron in the reporter into Neuro-2a cells (N2a) and observed strongly correlated red and green fluorescence using flow cytometry (Fig. 5b), as expected. To test the effects of RNA structure across this 5’SS, a 37-nt insert was designed and inserted downstream to generate a 22-nt stem loop that sequesters upstream sequence including the 5’SS (Fig. 5c). We also designed and tested single and double intronic point mutants to increase stem loop EFE, as well as a fully destabilized and an unstructured construct where base pairing with surrounding sequence is unfavorable (Fig. 5c). Importantly, the point mutations are not directly opposite the 5’SS; based on the results of our experiments in yeast (see Fig. 4), we expected an increase in fluorescence due to decreased overall stem loop stability and thereby increased access to the 5’SS. Indeed, the perfect stem loop drastically decreased gene output compared to the wildtype *HBB* intron; this decrease was slightly rescued with a single point mutant (Fig. 5d). Analysis of all constructs in both N2a and HeLa cells enabled us to calculate gene output from the log ratio of mNeon and mScar fluorescence (Fig. 5e). Strikingly, a stepwise increase in gene output was observed as the stability of the stem loop decreased in both cell lines. Fully unstructured inserts outperformed the structured wildtype sequence, and splicing efficiencies in the transfected cell populations were confirmed by RT-PCR (Fig. S5b). Taken together, these results confirm that fine-tuned regulation of splicing through RNA structure formation is a conserved mechanism in higher eukaryotes.

**Fig. 5:**
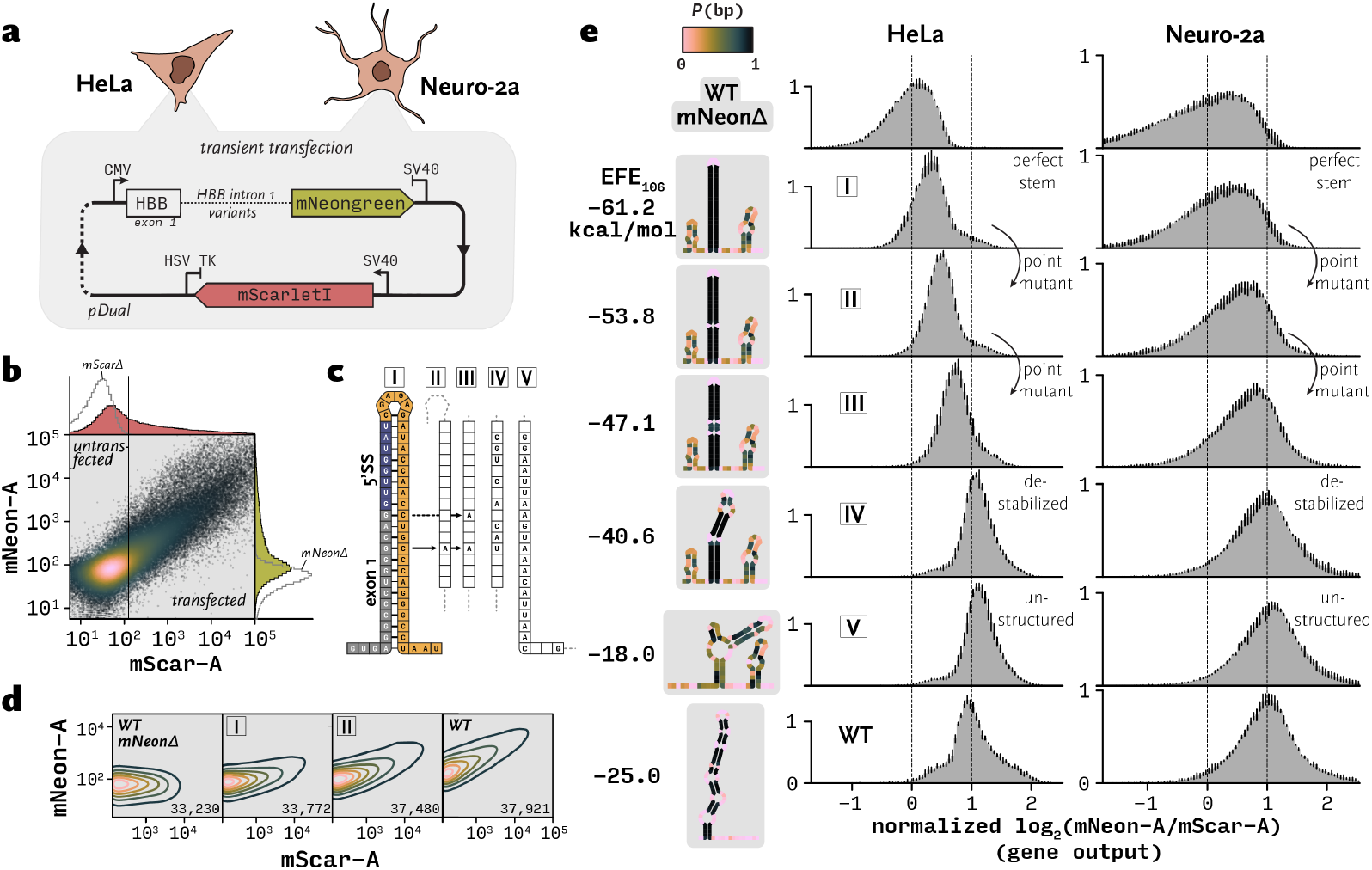
Stem loop stability across 5’SS regulates gene output in mammalian cells. **(a)** Dual expression plasmid for transfection into tissue culture cells. **(b)** Flow cytometry of N2a cells transfected with pDual carrying the wildtype *HBB* intron. Transfected cells are determined using an mScar fluorescence cutoff chosen based on a control with truncated, non-fluorescent mScar (*mScarΔ*). **(c)** Intron designed to sequester the 5’SS in a perfect stem loop (I) and destabilized versions of I, including single (II) and double (III) point mutants, fully destabilized (IV) and unstructured (V) variants. **(d)** Flow cytometry measurements of mScar and mNeon fluorescence of N2a cells transfected with the indicated constructs (only transfected cells shown). Number of transfected cells is indicated at the bottom. **(e)** Gene output for both HeLa and N2a cell lines for each construct. Median gene output of the wildtype *HBB* intron construct with truncated mNeon (*mNeonΔ*) was set to zero and the wildtype *HBB* intron construct with intact mNeon was set to one. Error bars indicate mean and standard deviation from three biological replicates. MFE structures with overlayed pairing probabilities are shown to the left. EFE is indicated.

### Structure-destabilizing variants emerge rapidly under selective pressure

If intron sequence is relatively malleable, and point mutations are enough to destabilize an inhibitory structure and increase gene output detectably, we would expect to see intronic mutations emerge when a cell population faces selective pressure against a stable intronic RNA structure. To test this, we attached a KanMX cassette to mNeon in the yeast system, which allowed us to apply selective pressure by adding the antibiotic G418 to growth media. We placed a stable stem loop across the 5’SS, chosen based on its low fluorescence in the Sort-seq experiment. We used an intronless mNeon-KanMX strain as a control. Since gene expression was not completely inhibited by the stem loop, the resulting yeast strain grew on moderate concentrations of G418 (permissive, 200 µg/ml), but was inviable at high concentrations (restrictive, 1.5 mg/ml), while the intronless strain grew at both concentrations. We subjected cultures of both strains to UV irradiation to induce mutations in random locations throughout the genome and plated the yeast on restrictive or permissive G418-containing plates. Both strains form lawns on permissive plates; on restrictive plates, the intronless strain forms a lawn and the strain with a structured intron forms colonies, of which we isolated a pool of around 1,000. We isolated genomic DNA from all plates and sequenced most of exon 1 and the entire intron region to calculate mutation rates (Fig. 6a). As expected, mutations emerged only on restrictive plates only for the intron-containing strain, and not in any of the controls. Most of these mutations were in the intronic region (Fig. 6b). The most abundant mutant changes a U-A into a U·G pair and thus marginally destabilizes the stem that sequesters the 5’SS; evidently enough to increase gene output above the survival threshold (Fig. 6c). Another notable mutant changes the optimal GAGA (GNRA) tetraloop into GAGG, which is expected to decrease loop stability. This subtle change in tertiary structure granting survival highlights the important role that RNA structure plays in gene expression. Indeed, almost all mutants we isolated decreased the predicted EFE of the stem loop through intronic mutations (Fig. 6d). These results demonstrate that subtle changes to secondary or tertiary RNA structure, caused by point mutations, can modulate gene expression enough to alter cellular survival.

**Fig. 6:**
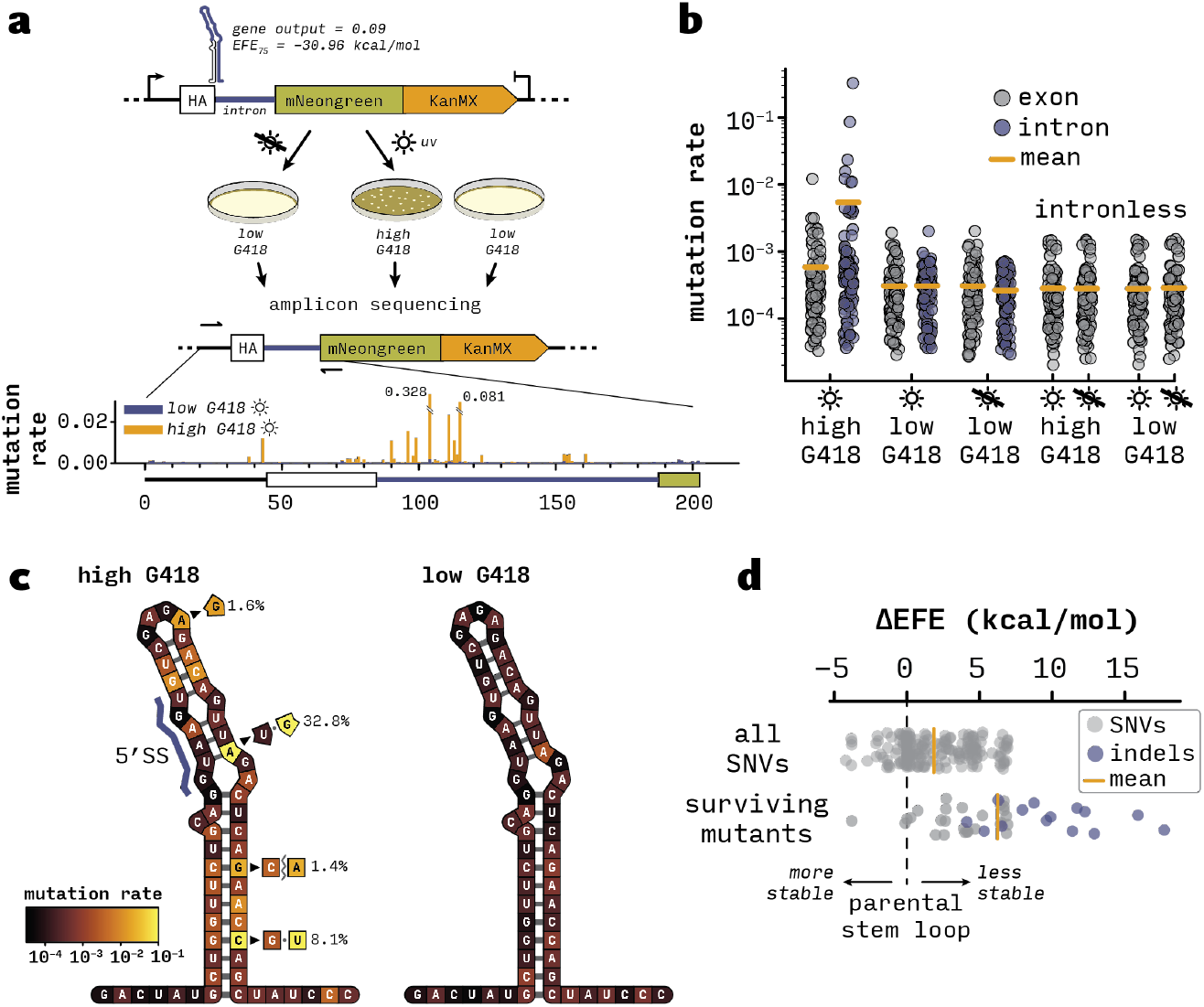
Under selective pressure, intronic RNA structure disrupting mutations emerge. **(a)** Experimental scheme to enrich yeast strains with mutations that increase gene output. Selective pressure is applied by attaching a G418-resistance cassette to the fluorescent splicing reporter. The parental strain harbors a poorly splicing intron with a relatively stable stem loop structure across the exon-intron junction. Treatment with UV light followed by selection on permissive (low concentration) or restrictive (high concentration) G418 plates leads to a yeast lawn or single surviving colonies, respectively. The locus around the intron is amplified from gDNA and sequenced. Mutation rates are calculated at every position of the amplicon. **(b)** Mutation rates for intron and exon regions under different selection conditions. **(c)** Mutation rates overlaid on the predicted MFE structure of the parental exon-intron junction for yeast grown on restrictive (left) or permissive (right) G418 plates. Allele frequencies are given in percent. **(d)** Predicted EFE difference compared to the parental exon-intron junction. All possible SNVs define the background set. Surviving mutants were isolated from restrictive G418 plates with UV treatment.

## Discussion

The biological function(s) of introns have long been debated. By normalizing genetic background and changing only intron RNA structure in our MPRA, we systematically reveal the striking relationship between RNA structure and gene output by predictably tuning the efficiency of pre-mRNA splicing. As proof of principle, we were able to alter human β-globin gene output by designing structured 5’SSs with a range of stabilities following parallel design rules. This substantiates the idea that introns, which are less constrained in their sequence than exons, may exhibit widespread changes in pre-mRNA folding that have been selected to benefit the organism. Conversely, intron sequences and structures can be causative in the context of disease-associated intronic mutations (*11, 45*). Our findings thus suggest ways to therapeutically address harmful variants.

Unexpectedly, we discovered a region of the intron that is sensitive to structure formation without involving the splice sites; in these cases, the presence of a stem loop upstream of the branch site decreased gene output. A plausible explanation is that recruitment of the U2 snRNP may be sterically hindered by these stem loops during branch site recognition, which may be owing to constraints imposed by U2 snRNP proteins (*46*). We speculate that structures in these intron regions could be evolutionarily selected for to dampen gene expression. Integrated controls in our library design show that the functional disruption is not due to sequence change but relies solely on structure. This method of scanning a functional sequence for sensitivity to base pairing may be applicable to several other systems in future.

The 5’SS and branch site are recognized by RNA-RNA intramolecular base pairing with spliceosomal snRNAs U1 and U2, respectively. Note that the relative insensitivity to structures engaging the 3’SS likely reflects the activity of the spliceosome as an unwindase that can find the optimal CAG nucleotides in the transition between the first and the second steps of catalysis (*47*). A strong negative correlation between structure stability and gene output at 5’SSs and branch sites suggests that intramolecular base pairing directly competes with snRNA annealing. Since intramolecular base pairing occurs rapidly (*4*), recruitment of either U1 or U2 snRNP must in these cases be promoted by other processes, such as exon definition or intron cross-bridging interactions. Specific association of U1 or U2 snRNPs before intermolecular base pairing takes place might then enable a strand invasion process that would in turn depend on stem loop stability, consistent with our data. This model suggests that – in the face of competition from intramolecular base pairing with their target splice sites – the U1 snRNP may be recruited as a consequence of an engaged U2 snRNP, or vice versa. Single molecule measurements indicate the order of U1 and U2 recruitment is flexible, though RNA folding during the experiment was not specifically considered (*48*).

Taken together, our findings indicate that RNA folding is a selectable feature of introns, much as transcription factor binding sites are selectable features that can change promoter activity. Consistent with this view, we show that when applying selection pressure against a 5’SS-sequestering intron structure, destabilizing intronic mutations that permit survival emerge (see Fig. 6). In contrast to other sequence elements, numerous RNA structure configurations may achieve the same regulatory effect, likely weakening covariation signatures that we typically consult to detect functional elements (*49, 50*). These structures would be most effective if they form co-transcriptionally, since many introns are removed while the nascent transcript is being elongated by Pol II; those that are not removed co-transcriptionally could also undergo regulation by structure after 3’-end cleavage with the potential involvement of many RNA binding proteins and helicases in cells (*5, 6*). Our study shows that machine learning, including the application of neural networks, predicts gene output well in our well-defined system, suggesting that algorithms predicting splicing behavior would be improved by the inclusion of intron structure information (*51, 52*). However, a significant current obstacle is predicting or measuring the RNA folding landscape of all introns (*10*). Impactful structures can include both intron and exon sequences as we have seen for β-globin. These regions may be most predictive of splicing in the near future and enable further exploration of thus far enigmatic disease-associated human genotypes.

## Acknowledgements

The authors would like to thank Srikar Krishna Gopinath, Jernej Ule and Manny Ares for helpful discussions and Sabrina Hu and Leena Sen for assistance with library cloning. This work was supported by the National Institutes of Health R01GM112766 (to KMN) and a predoctoral fellowship from the American Heart Association (908949 to LS). Data acquisition at Yale Center for Genomic Analysis was supported by the National Institute of General Medical Sciences of the National Institutes of Health under Award Number 1S10OD030363-01A1. This work is solely the responsibility of the authors and does not necessarily represent the official views of the NIH. **Author contributions:** Conceptualization, L.S.; Methodology, L.S.; Investigation, L.S., P.B., and P.P.-B.; Visualization, L.S.; Software, L.S.; Writing – Original Draft, L.S. and K.M.N.; Writing – Review & Editing, L.S. and K.M.N; Supervision, K.M.N. and L.S.; Funding Acquisition, K.M.N. and L.S. **Data and materials availability:** Raw sequencing data are available through the National Center for Biotechnology Information (NCBI) under GEO accession number GSE308590. Flow cytometry FCS files are available at https://doi.org/10.5281/zenodo.17160682. Code is available at https://github.com/lschaerfen/intron_sortseq. Materials are available upon request. **Competing Interests:** The authors declare no competing interests.

## Supplementary Materials

Materials and Methods

Figs. S1 to S5

Table S1

## Supplementary Materials

## Methods

### Strain construction

To allow for fast and efficient integration of intron variants into the *S. cerevisiae* genome, we constructed a receiver strain stably expressing mScarletI, inducibly expressing Cre recombinase, and harboring a *lox66*/*splitHygR* landing pad. We integrated red fluorescent mScarletI driven by *TDH3* promoter and *CYC1* terminator at the *HIS3* locus using homologous recombination and selection on synthetic complete dropout media lacking histidine. The landing pad constitutes the 5’-half of the *hphMX3* hygromycin resistance cassette and half of a synthetic intron containing a *lox66* site, which was integrated at the *MET17* locus using homologous recombination and selection on synthetic complete dropout media lacking methionine. We then transformed the resulting strain with a plasmid expressing Cre recombinase from the galactose inducible *GAL10* promoter and selection on synthetic complete dropout media lacking uracil. The resulting strain BY4741 *his3::mScarletI met17::land66 pGAL-URA2-Cre* is competent for site-specific integration of transgenes delivered *via* the payload plasmid *pPL*. This plasmid, for propagation only in *E. coli*, features a *lox71* site followed directly by the 3’-half of the synthetic intron and the 3’-half of hphMX3. The payload constitutes an HA-mNeongreen expression cassette driven by *TDH3* promoter and *CYC1* terminator interrupted by a Golden Gate cassette harboring self-excising SapI/BspQI type-II restriction sites for efficient intron library insertion.

### Yeast library construction

Intron libraries were ordered as single-stranded DNA oligonucleotides flanked with primer binding regions containing SapI/BspQI sites. Libraries were amplified for 10 cycles by PCR with KAPA HiFi HotStart polymerase (Roche) and purified with 1.3X AMPure XP beads (Beckman). Golden Gate reactions were carried out with 3 nM pPL plasmid and 6 nM purified intron library, 5 U/µl T4 DNA ligase (NEB) and 0.75 U/µl SapI (NEB) and 1X T4 DNA Ligase reaction Buffer (NEB). Reactions were cycled 40 times for [1 min at 37 ºC, 1 min at 15 ºC], then incubated 5 min at 37 ºC, 5 min at 60 ºC and then transformed into Stellar (Takara) competent *E. coli* cells. Depending on library size, multiple transformations were combined and transformants were incubated over night at 37 ºC in 500 ml LB media supplemented with 100 µg/ml carbenicillin. Plasmid DNA was extracted from *E. coli* libraries with the QIAGEN® Plasmid Midi kit. Next, *S. cerevisiae* BY4741 *his3::mScarletI met17::land71 pGAL-URA2-Cre* was transformed with plasmid library DNA using the lithium acetate method: yeast cultures were grown in synthetic complete dropout media lacking uracil until exponential phase (OD_600_ around 0.6-0.7). 9 OD_600_ units were harvested and washed with 100 mM LiOAc, then resuspended in 100 µl 100 mM LiOAc. 100 µg sheared, denatured Salmon Sperm DNA (Invitrogen) and 5 µg plasmid library were added to a final volume of 210 µl and incubated at room temperature for 30 min. A mixture comprising 100 µl DMSO, 90 µl 1 M LiOAc, 600 µl 50% PEG 3350 was added, and the reaction was incubated for 30 min at room temperature. Next, transformation reactions were incubated for 15 min at 42 ºC, then pelleted, and then resuspended in 250 µl 4 mM CaCl_2_ and incubated for 10 min. Each transformation was plated onto two YPAG plates (10 g/l yeast extract, 20 g/l peptone, 20 g/l agar, 40 mg/l adenine hemisulfate, 2% (w/v) galactose) to induce Cre expression and incubated at 30 ºC for 24 h. Transformants were then replica plated onto selective YPAD (YPA with 2% glucose and 200 µg/ml hygromycin-B) plates and incubated for 36 h. Colonies were counted, requiring at least *N*(log(*N*) + *γ*) colonies for a library of size *N*, where *γ* is the Euler-Mascheroni constant. Colonies were then scraped off the plates and diluted in 500 ml YPAD with 200 µg/ml hygromycin-B to a starting OD_600_ of 0.05. The resulting yeast library was grown to stationary phase and stored at -80 ºC in YPA with 15% glycerol.

### Tissue culture and transfection

Neuro-2a cells or HeLa cells were cultured in DMEM media with Glutamax (Gibco #10569-010), supplemented with 100 U/ml Penicillin, 100 µg/ml Streptomycin, and 10% fetal bovine serum. 2×10^5^ cells/well were seeded in a 6-well plate 24 h before transfection with Lipofectamine 3000 (Thermo Scientific) and 2 µg of pDual plasmid following the manufacturer’s protocol.

### Cell Sorting and Flow cytometry

Yeast libraries were grown in liquid YPAD to exponential phase (OD_600_ around 0.6-0.7), harvested and washed with ice cold 100 mM potassium phosphate buffer (PPB), then resuspended in 100 mM ice cold PPB for subsequent sorting. Cells were sorted into approximately evenly spaced bins of mNeongreen over mScarletI fluorescence emission ratio (488 nm and 635 nm excitation, respectively) using a BD FACS Aria III sorter. Fresh YPAD cultures were started from the sorted populations, grown over night, and each population was again sorted into four bins, yielding 16 cultures in total. Sorted populations were grown overnight and the output fluorescence distributions were acquired by flow cytometry using the same settings (Tab. S1). HeLa and Neuro-2a cells were analyzed 18 h after transfection with a BD LSR Fortessa cytometer. After removal of doublets, cells with mScar fluorescence below or equal to the non-fluorescent, truncated (*mScarΔ*) control were removed. Fluorescence compensation parameters were set for each fluorophore. For yeast flow cytometry, doublet discrimination or compensation were not needed. Gene output for both species was calculated as the log2 fraction of mNeon over mScar fluorescence.

### High-throughput amplicon sequencing

For each bin and for the input library, variant intron sequences were amplified from gDNA using PCR. Yeast cells were harvested and resuspended at low density in H_2_O. Cell walls were digested in 20 µl reactions with 1 µl Zymolase (Zymo Research) in 25 mM sodium phosphate pH 7.5 for 20 min at room temperature, then 5 min at 37 ºC. Samples were diluted with 60 µl H_2_O and 2 µl of the resulting crude gDNA was used as template for 15 cycles of PCR with KAPA HiFi HotStart polymerase (Roche) with primers adding Illumina flowcell binding sequences. Reactions were purified using AMPure XP beads (Beckman) and used as template for a second round of PCR with Illumina TruSeq i5 and i7 index primers for 15 cycles. Final libraries were again purified with AMPure XP beads, pooled and sequenced on Illumina NovaSeq machines in 2×150 paired-end mode.

### Yeast selection experiments

For selection experiments, we first modified the payload plasmid pPL by appending a G418 resistance gene (*Kan*^R^) to the mNeongreen coding sequence, yielding pPL-Kan. We then integrated the intronless version of pPL-Kan and a version with a *QCR10* intron variant with a relatively stable stem across the 5’SS. These strains were grown to exponential phase and 10 OD units were irradiated in 10 ml H_2_O in 15 cm petri dishes with 254 nm UV light at a dosage of 30 µJ/mm^2^. After irradiation, cells were incubated at 30 ºC in YPAD for 1 h and then plated on two permissive YPAD + 200 µg/ml G418 plates, incubated at 30 ºC over night, and then replica plated onto either restrictive YPAD + 1.5 mg/ml G418 or onto permissive plates as a control, and then grown for two days. Yeast colonies or lawns were scraped off the plates and genomic DNA was isolated. Amplicon sequencing was performed as described above. Paired-end reads were pre-processed and merged with fastp (*1*) and then aligned to the respective reporter gene sequence using BWA (*2*). Positional mutation rates were calculated from pileups generated by samtools mpileup. The nature and frequency of individual mutation events was obtained using LoFreq (*3*).

### Variant identification

Reads for all bins and for the input library were first processed with fastp, (*1*) merging read pairs into a single read. For intron libraries with known sequence, the resulting fastq files were parsed directly and reads were counted if the sequence was matching a variant in the library perfectly. This naïve counting method assigned 70-90% of reads to a unique variant.

For randomized libraries, intron variants are unknown. We therefore first concatenated fastq files for all bins and the input, then extracted read sequences and stored those in a text file. This file was then sorted using the Linux “sort” command, and unique sequences and their counts were extracted using the Linux “uniq” command. Sequences that had less than *x* counts (*c*) were then removed by empirically setting *x* to the position of the shoulder of the survival function

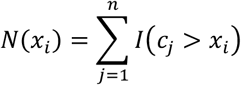

where *n* denotes the total number of sequences and *I*(•) is the indicator function. Sequences that passed this filter were then clustered using the Clusterize algorithm (*4*) with a similarity cutoff value such that the number of sequences inside clusters remained low. For each cluster, the consensus sequence was determined and retained. Sequences not in clusters were retained as well, and the total number of resulting sequence variants corresponded approximately to the number of yeast colonies obtained after library transformation. Reads for all bins were then assigned to these variants using the naïve approach described above.

### Sequence Design

To design “walking stem” intron libraries, we wrote an automated pipeline that runs antaRNA (*5*) to design the 19-nucleotide inserts. First, for every insert position, we generated a set of inserts that, together with sequestered intron sequence, fold into the target stem loop structure. The target structure is a 13-nucleotide stem capped by a tetraloop with two nucleotides each upstream and downstream of the stem’s base, of which the one closer to the base is designed to be unpaired, and the further one left unconstrained. The tetraloop and either side of the stem, as well as the two nucleotides adjacent to the stem comprise the 19-nt insert that is variable during the design process. From the antaRNA designs, we selected two upstream/downstream sequestering sequences each, ensuring a minimum Hamming distance of 3. The sequestering sequences were then input into the rf_mutate function from RNAFramework (*6*) to select two inserts harboring three point mutations that maximally destabilize the stem loop. We also selected ten 19-nucleotide inserts per position that were designed to leave the surrounding intron fully unpaired. All generated sequences were inserted into the original wildtype *GOT1* intron sequence, and primer binding sites for amplification and Golden Gate cloning were appended. The resulting library was ordered as an oligo pool from Twist Bioscience.

### Massively parallel fluorescence quantification

To assign an accurate fluorescence estimate to each variant, we first determined the median fluorescence value of the output distributions (*i*.*e*. the post-sorting distribution for each bin). These were scaled such that the negative control was 0 and the positive control was 1. For each bin separately, variant counts were normalized by multiplying with the number of variants and dividing by the sum of all counts. Counts for each bin were then scaled according to the number of variants found in that bin, which was determined by thresholding normalized counts with an empirically determined cutoff that depends on sequencing depth. The cutoff value is chosen by assessing pairwise Spearman correlation matrices for counts from all bins. For each variant, normalized, scaled counts are denoted as *y* and bin position (median of output fluorescence distribution) is denoted as *x*. A Gaussian distribution was then fitted to the resulting histogram using maximum likelihood estimation (MLE) with a negative log likelihood function

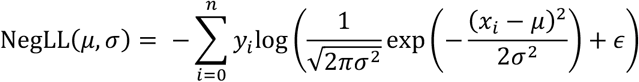

where *μ* and *σ* are mean and standard deviation of the Gaussian, respectively, *n* is the number of data points (usually 16) and ϵ is a small number to avoid the logarithm of 0. We used the LM-BFGS algorithm implemented in the “minimize” function from SciPy to find parameters that minimize NegLL(*μ, σ*), yielding estimated parameters 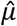 and 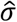 for each variant.

### Position-resolved structure impact

To systematically assess the effect of RNA structure formation across an intron, we calculated position-resolved summary statistics and structure impact scores across the *GOT1* intron sequence. Sequence variants belonging to stem-forming, destabilized (upstream/downstream), and control unstructured libraries were grouped by their annotated insertion positions within the intron. For each nucleotide position, fluorescence estimates *μ* from all variants predicted to sequester that position were aggregated and the mean fluorescence value was calculated. This approach measures the average phenotypic impact of forming a stem at each position, independent of the variant’s precise insertion origin. For each position, we then computed a structure impact score as follows:

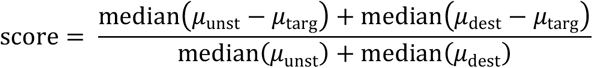

where *μ*_targ_, *μ*_dest_, and *μ*_unst_ denote the fluorescence values for, respectively, stem-forming, destabilized, and unstructured variant sets sequestering that position.

### Structure prediction

All structure prediction was carried out using the ViennaRNA (*7*) Python API. The sequence length for structure prediction was optimized as shown in Fig. 2a and Fig. S3a. The temperature was set to 30 ºC. First, the partition function was calculated and ensemble free energy, positional entropy, ensemble diversity and pairing probability matrix determined. The matrix was summed to obtain the *P*(bp) vector. RNA structures were visualized using coordinates calculated with the RNApuzzler algorithm (*8*).

### Machine learning

Random forest regression models made up of 200 decision trees were trained using the scikit-learn package for Python. Pairing probability vectors were used as predictors for splicing efficiency, and 85% of the data were used for training. The coefficient of determination *R*^2^ and its standard deviation were calculated using 30-fold cross validation. Neural networks with genotype-phenotype maps were trained using MAVE-NN (*9*). Only variable nucleotides were used as features for training, and data were randomly split into 80% training, 10% validation and 10% test sets. Using MAVE-NN, global epistasis regression models with homoskedastic Gaussian noise and pairwise interactions were trained for 750 epochs with a learning rate of 0.0005 and batch size of 50.

---

**Fig. S1:**
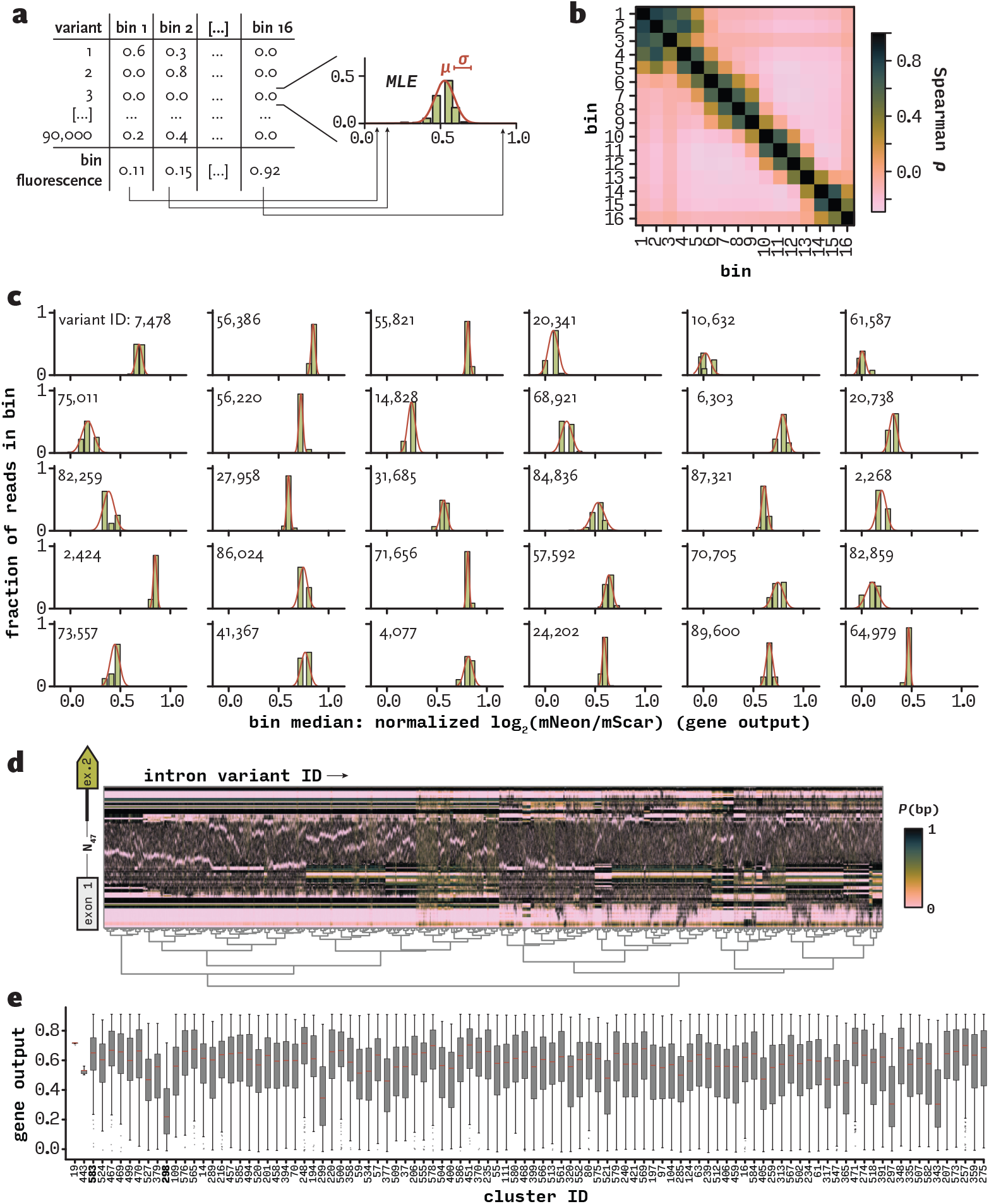
Robust fluorescence measurement for thousands of variants in parallel. **(a)** Principle of fluorescence estimation based on DNA sequencing data from each bin. The fraction of total reads for each variant occurring in each bin is calculated from Illumina sequencing data. These fractions represent histogram frequencies with bins positioned at the empirically determined median fluorescence value for each bin. Maximum likelihood estimation (MLE) is then used to infer the original fluorescence value *µ* and its standard deviation *σ*. **(b)** Pairwise Spearman correlation coefficients of read counts for all variants between bins. **(c)** Normalized read count frequency distributions and MLE fits for randomly selected variants. **(d)** Hierarchical clustering of *P*(bp) vectors. Each column represents one variant; each row represents one nucleotide. **(e)** Gene output distributions for variants grouped by hierarchical clustering. Sorted by standard deviation.

**Fig. S2:**
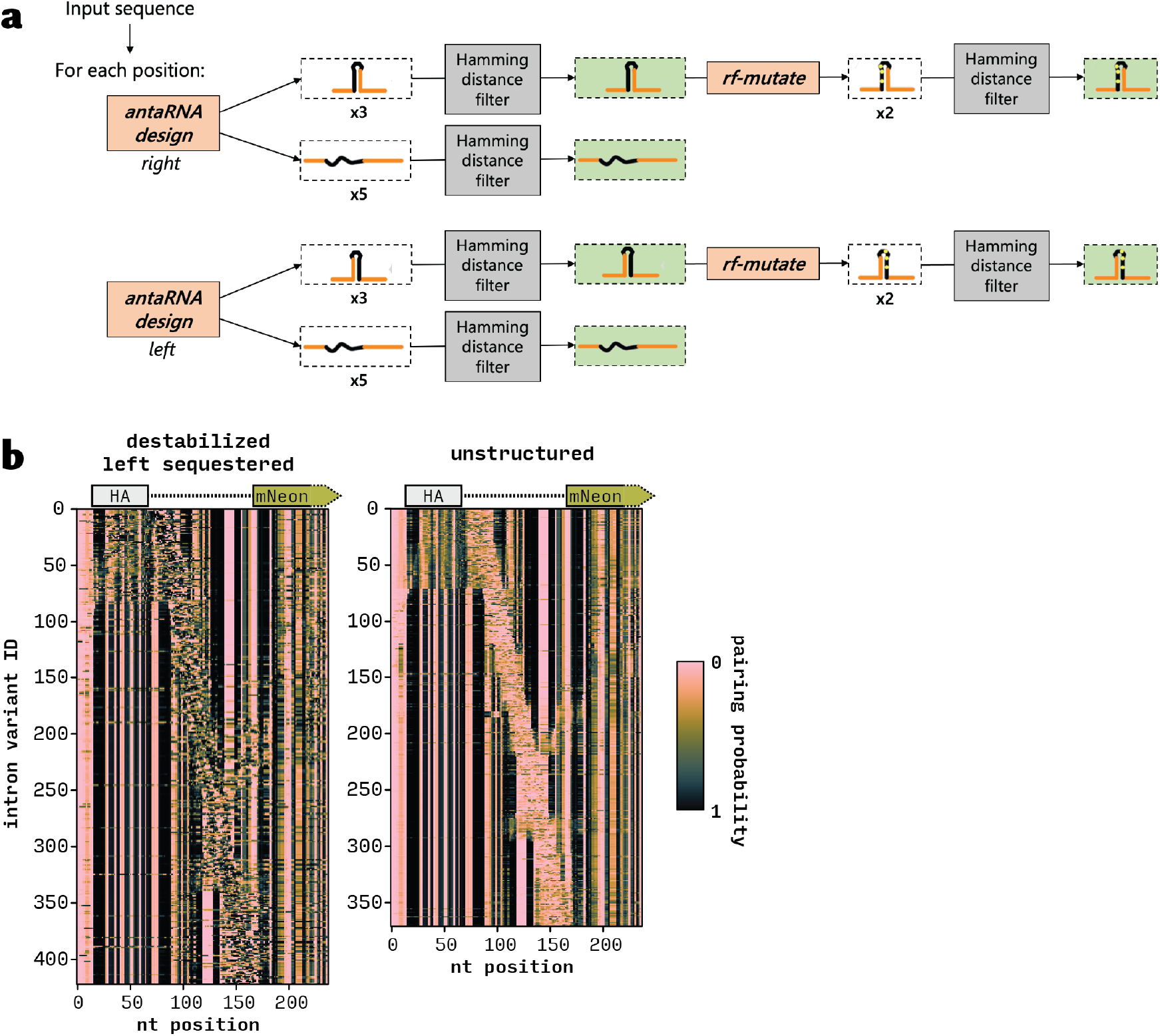
Structure-aware sequence design allows detection of structure-sensitive sequence elements. **(a)** Algorithmic design of 19 nt insert sequences to form upstream or downstream sequestering base pairing interactions, as well as destabilizing and unstructured controls. **(b)** Predicted *P*(bp) for each nucleotide of the designed *GOT1* intron variants harboring the destabilizing mutations in the insert sequestering upstream sequence (left) and the unstructured inserts (right). Each row of the matrix represents one unique intron.

**Fig. S3:**
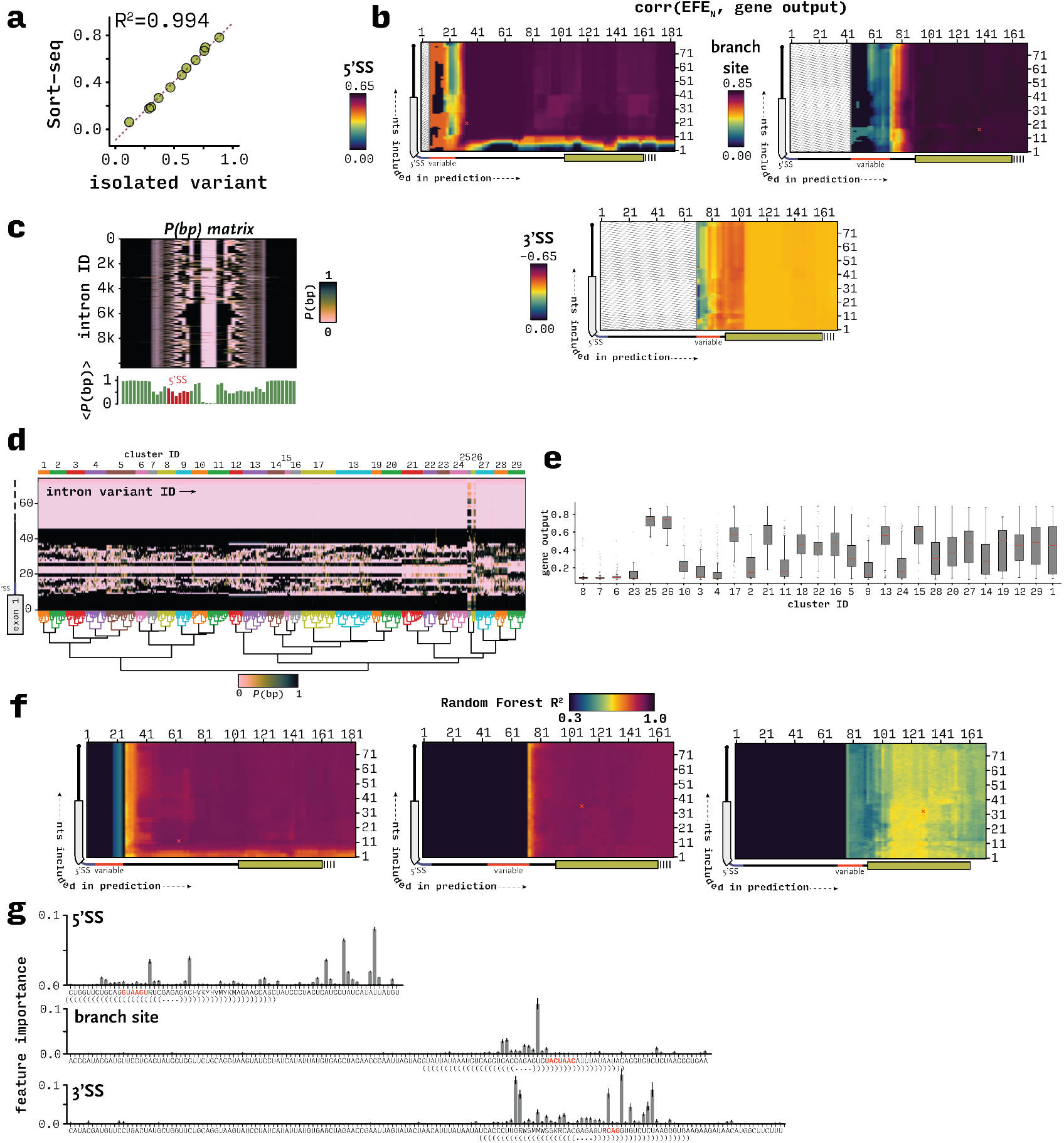
RNA ensemble structure prediction, clustering and machine learning. **(a)** Comparison of gene output measurements obtained from flow cytometry of individually isolated variants, and from Sort-seq. Variants are from the 5’SS library. **(b)** Pearson correlation coefficients of EFE and gene output for varying structure prediction windows. Y-axis represents the start coordinate of the window (number of nucleotides included upstream of 5’SS); x-axis represents the end coordinate (number of nucleotides included downstream of 5’SS). **(c)** *P*(bp) matrix for the 5’SS library. Bar plot below shows the positional average. **(d)** Hierarchical clustering of *P*(bp) vectors for the 5’SS library. Each column represents one variant; each row represents one nucleotide. **(e)** Gene output distributions for 5’SS library variants grouped by hierarchical clustering. Sorted by standard deviation. **(f)** Coefficient of determination for random forest regressors. Axes are the same as (b). **(g)** Feature importance from random forest regressor models trained on *P*(bp) to predict gene output.

**Fig. S4:**
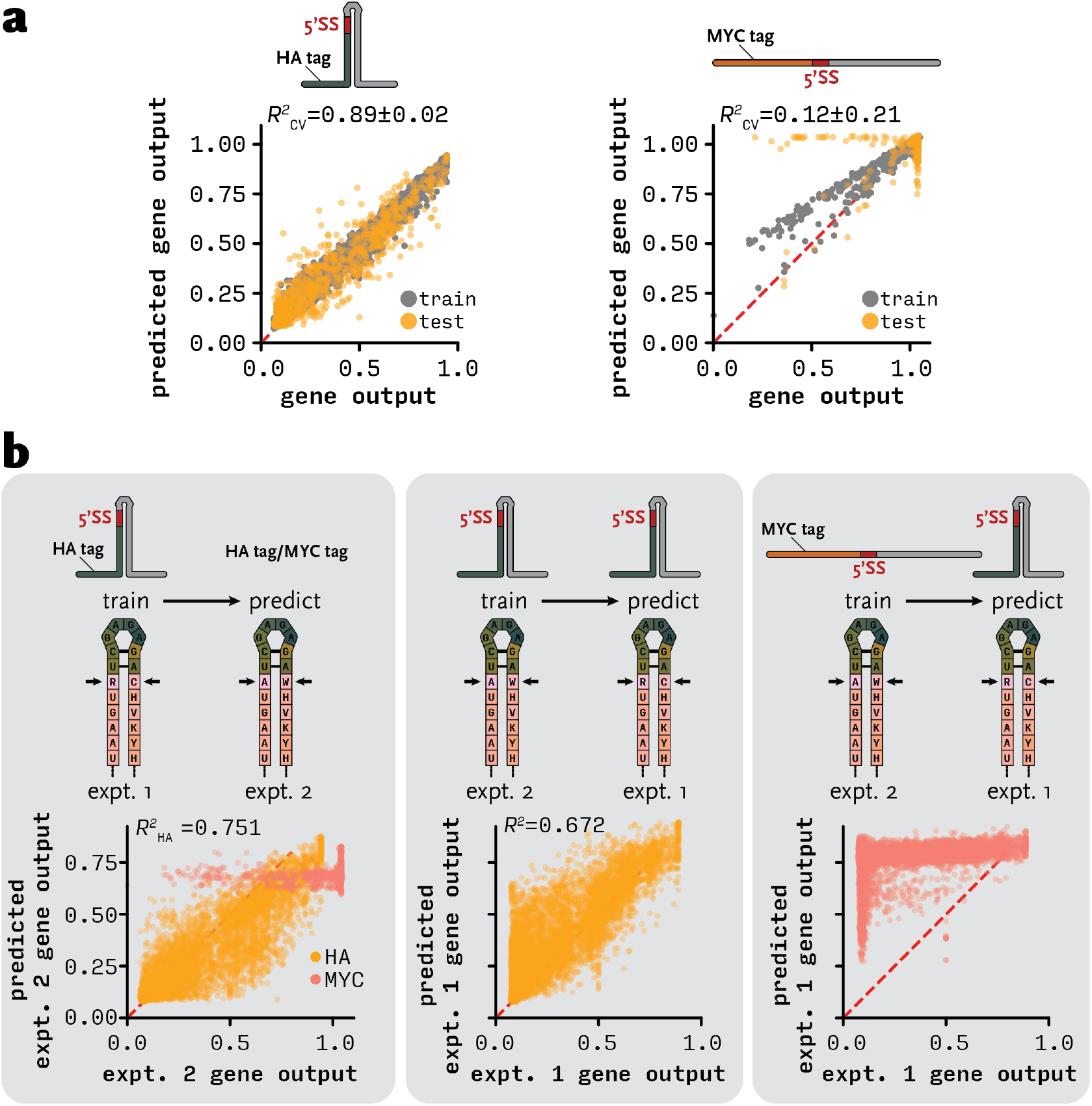
Reproducibility and sequence-independence of gene output predictions. **(a)** Performance of random forest regressor models trained to predict gene output from *P*(bp) for a second set of 5’SS sequestering variants (see Fig. 4b alternative bases). This second library was cloned with HA or MYC as exon 1 (same intron variants). **(b)** Random forest regressor prediction performance across separate experiments and libraries.

**Fig. S5:**
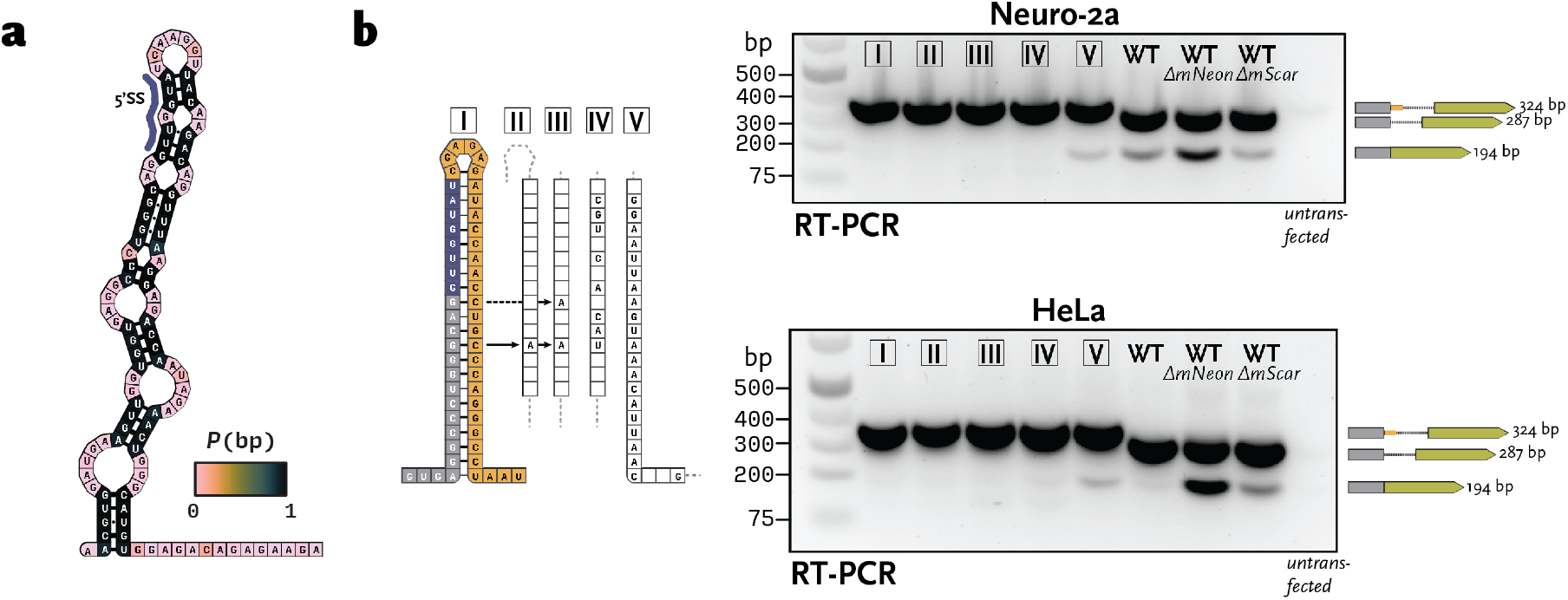
Regulation of mammalian pre-mRNA splicing by a sequestered 5’SS in human β-globin. **(a)** Predicted minimum free energy structure of exon 1 – intron junction of the wild-type (WT) human HBB gene. Colored by *P*(bp). **(b)** Agarose gels showing reverse transcription – PCR (RT-PCR) products with primers situated in exon 1 and exon 2 of the pDual construct.

**Tab. S1:**
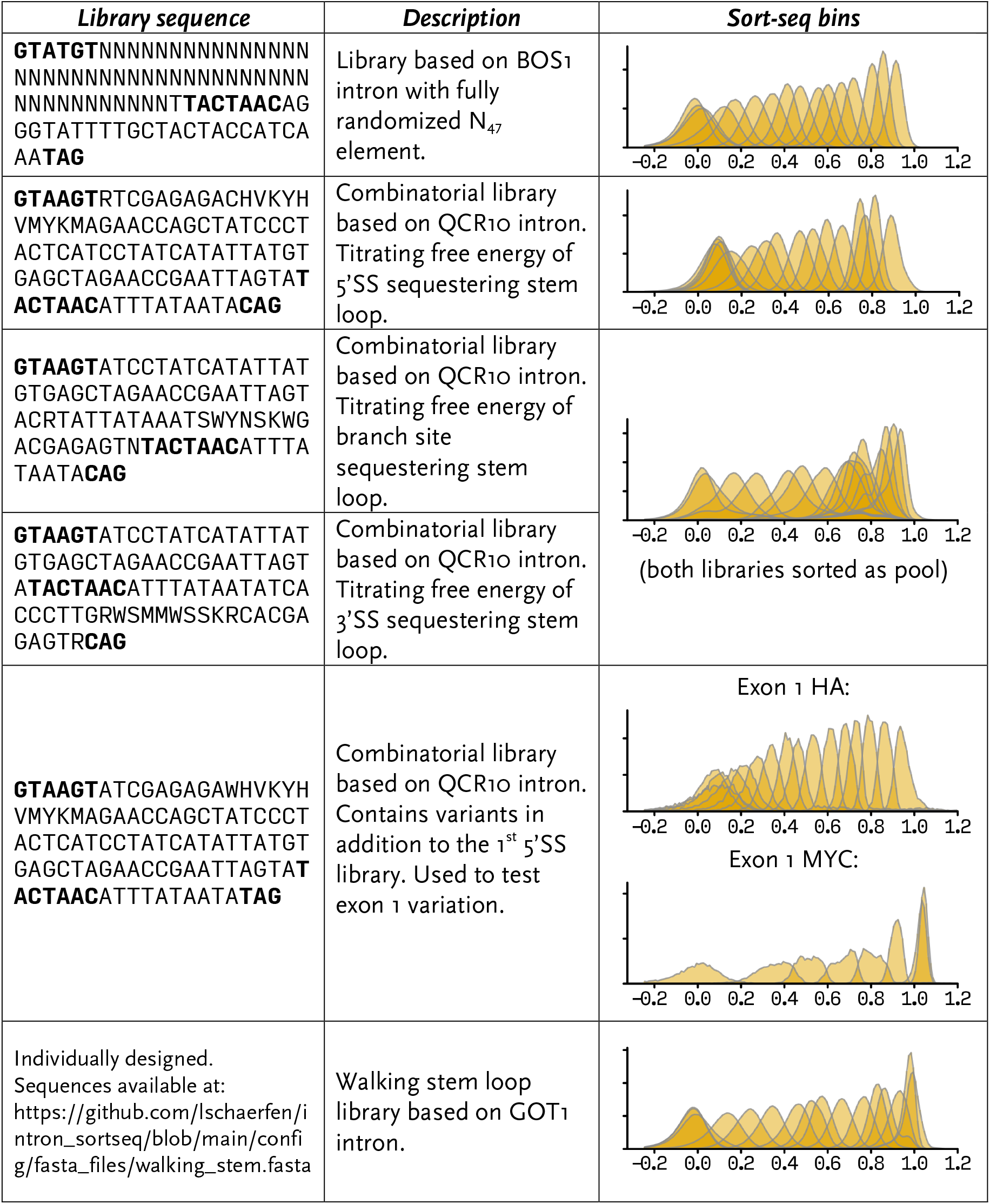
Library sequences and fluorescence output distributions. Gene output distributions for each individual bin were measured by flow cytometry after cell sorting. Individual histograms are normalized by cell count.

